# Designing Rigid Protein Fiducials to Visualize GPCR Conformational States

**DOI:** 10.64898/2026.01.24.701540

**Authors:** Alina Vo, Yuan-En Sun, Justin G. English, Michael J. Robertson

## Abstract

G-protein coupled receptors (GPCRs) mediate precise ligand-specific signaling profiles, yet structural visualization of how ligands alter receptor conformational landscapes in the absence of signaling partners or mimetics has proven incredibly challenging. Here we show that by combining generative protein design with deep-learning based conformational ensemble prediction we can reliably design ‘fiducial markers’ to facilitate cryogenic electron microscopy (cryoEM) of GPCRs at arbitrary fusion points, enabling the visualization of previously intractable states. We validate the approach with high-throughput determination of inactive state structures of four pharmaceutically relevant GPCRs, allowing for key details of receptor pharmacology to be resolved in each case. We then engineered an extracellular fiducial marker for the prototypical β2-adrenergic receptor that enabled direct structural characterization of the rearrangement of key intracellular motifs in the absence of G-protein. Comparison with recent co-folding models highlights gaps in current methods for predicting ligand-induced GPCR conformational changes. These results present a generalizable framework for accessing traditionally inaccessible structural states of small, dynamic proteins.

## Introduction

CryoEM has emerged as a powerful tool for studying structures and conformational ensembles of difficult to crystallize proteins. However, technical limitations prevent applying the technique to small proteins, which has complicated the study of the full conformational space of GPCRs, one of the largest classes of drug targets^1^. Several families of GPCRs, including the most extensive (family A), are too small to obtain high resolution reconstructions for the receptor alone with current technology (and indeed fall under the proposed ‘theoretical size limit’^2^). Several hundred structures of fully activated receptors bound to G-protein have been obtained with cryoEM given the significantly larger complex, but these snapshots represent a single physiologically transient state which often provides little insight into how ligands obtain precise pharmacological profiles, including functional selectivity^3^ and varying levels of agonism. To enable visualization of additional conformational states, ‘fiducial markers’, rigid binders or fusion proteins that provide additional molecular weight, have been employed to facilitate particle alignment. However, in the case of GPCRs, fiducial markers almost exclusively bind or fuse to intracellular loop 3 (ICL3), masking the native conformation, biasing the receptor towards the inactive state, and requiring experimental optimization^4-6^.

A variety of recent studies performing spectroscopic techniques^7-11^ have suggested agonists of family A GPCRs induce a complex, multi-state conformational landscape with intermediates between the fully inactive state, with transmembrane helix 6 (TM6) swung in, and a fully activated state where TM6 is completely opened, the latter of which is proposed to largely occur only in the presence of G-protein (or mimetics). Further, these studies have demonstrated that ligands with different signaling profiles differentially shift the populations of individual substates. Design of extracellular, rigid, pharmacologically inert fiducial markers would facilitate a structural understanding of these substates and how their populations shift in response to full, partial, and ‘biased’ agonists in a way that is not possible with ICL fusions. This in turn would benefit structure-based drug discovery of compounds with precision signaling profiles, which is a promising route to safer drugs^12^.

Generative machine learning methods for protein design have become incredibly powerful, providing a ‘bespoke’ solution for the production of arbitrary proteins^13,14^, but a robust and computationally efficient method for predicting the degree of flexibility between target and fiducial is necessary for reliable fiducial design (Figure 1a). Testing whether the highest-scoring AlphaFold2 (AF2)^15^ predicted structure matches the design model has become a standard component of protein engineering pipelines, but is often insufficient for fiducial marker prediction without additional cryoEM screening^16^. Metrics like AF2 local-distance difference test (pLDDT) score^17,18^ were also found to be less reliable in our assessment on a panel of well-characterized fiducial markers. Molecular dynamics simulations have demonstrated good performance for rigidity filtering^4^, but are far too resource intensive for design pipelines. Recently, various approaches for subsampling the MSA component and/or enabling dropout layers for AF2 together with extended sampling has been shown to be able to generate multiple conformations and alternate states of proteins^19-22^. However, less attention has been paid to whether a well-converged ensemble under such AlphaFold2-based approaches correlates well with molecular rigidity in cryoEM. It is also unclear how well these approaches, particularly those reliant on MSA alterations, perform when a part of the protein is engineered.

**Figure 1:**
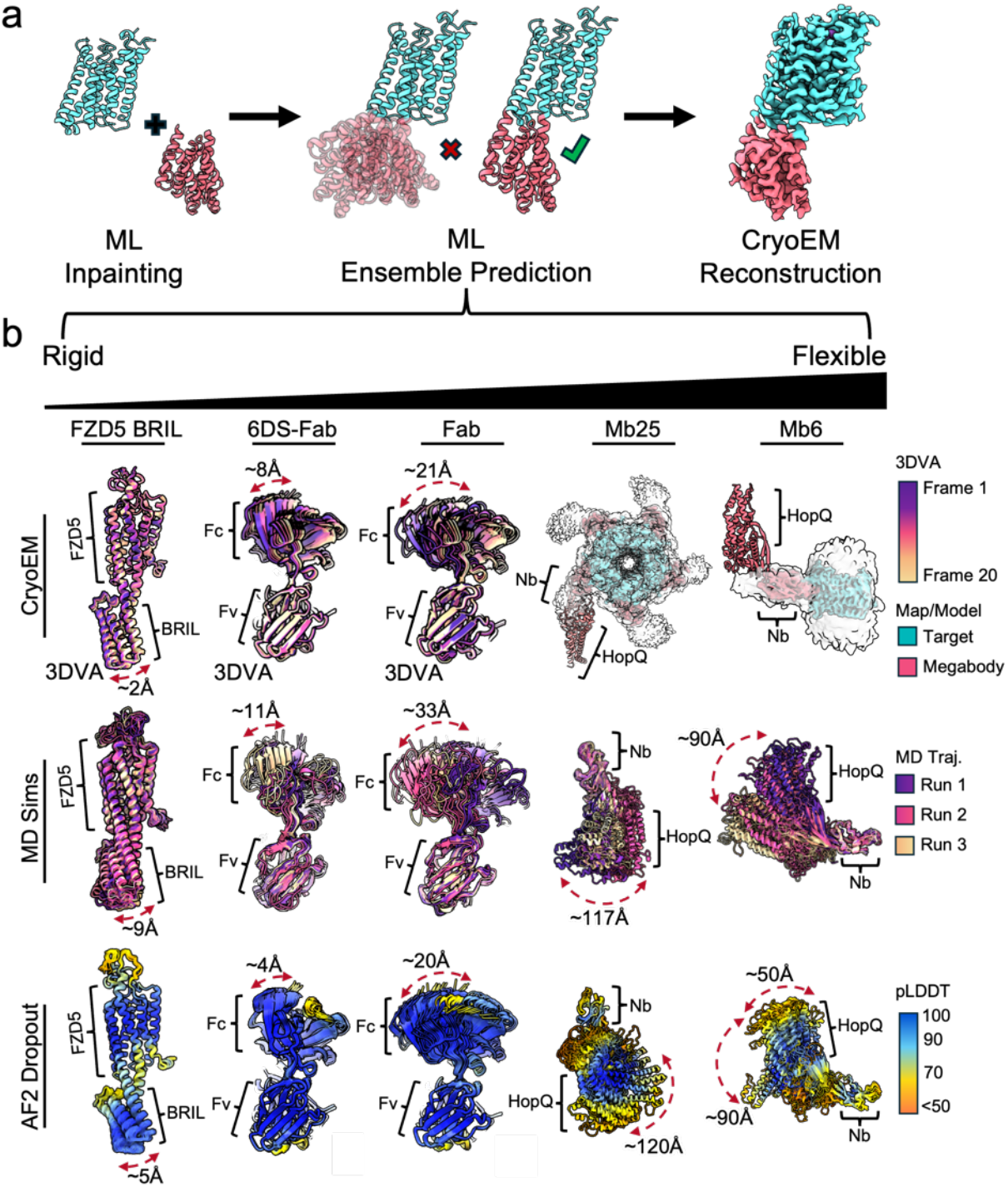
Predicting Fiducial Marker Rigidity. **a**, Schematic of the proposed design-rigidity filtering-structure determination pipeline. **b**, Comparison of published fiducial markers with raw cryoEM data deposited in EMPIAR. 3DVA was used to generate ensembles when the marker was sufficiently rigid; otherwise the reconstructed map is presented. Results are compared with equally spaced snapshots from triplicate 500 ns MD simulations (colored by replicate) and ensembles generated with AF2 with dropout enabled (colored by pLDDT score).

Here we demonstrate a correlation between the observed dynamics in cryoEM datasets of several popular fiducial markers (both binders and fusion partners), dynamics from MD simulations, and the predicted ensembles from AF2 extended sampling approaches^19-21^. We then leveraged AF2 sampling with dropout enabled for rigidity filtering together with generative design methods (RFDiffusion^13^ and ProteinMPNN^23^) to produce a rapid pipeline for obtaining inactive state structures of GPCRs based on ICL fusion, determining structures for four receptors bound to antagonists: the itch receptor MAS-related GPCR X2 (MRGPRX2), the adrenocorticotropic melanocortin receptor 2 (MC2R), vasopressin receptor 1A (V1AR), and gastric inhibitory polypeptide receptor (GIPR), an incretin receptor where both antagonism and agonism induce weight loss. This protocol was highly successful for predicting rigid ICL fusion constructs, with every design subjected to cryoEM analysis leading to a map of 3.4 Å resolution or better. These structures allow not only for a rationalization of antagonist binding, but also critical assessment of the predictive power of protein-ligand ‘co-folding’ models (OpenFold3 (OF3), Chai-1, Boltz-2^24-27^), which generally had issues with identifying GPCR-ligand poses and interactions. Finally, we designed a panel of extracellular fiducial markers for a well-studied GPCR, β2 adrenergic receptor, fused to TM1. The most successful design enabled structure determination in the presence of an antagonist, a strong agonist, a Gi-biased agonist, and complex with Gs protein. Supported by MD simulations, these results revealed an improved picture of the activation process and subtleties in the movement of key motifs that, together with prior spectroscopic results, emphasize the role of not just ordered TM5, TM6, and TM7 movement but also more dynamic motifs in the activation process.

## Results

### Benchmarking Prediction of Fiducial Rigidity

To assess methods for predicting the rigidity of the fusion between our GPCR targets and any designed proteins, we assembled a test set of fiducial markers that have been imaged in cryoEM (comprised of both fully native proteins and engineered) spanning from rigid to extremely flexible, including two example pairs where an attempt was made to rigidify a fiducial marker that was found to be too dynamic (the antibody Fab region rigidified with engineered disulfides^28^ and ‘macrobodies’ with engineered prolines^29^, Figure 1b, Extended Data Figure 1a,b). These sets were grouped into cases where raw micrographs were available in the EMPIAR database, which allowed for reprocessing the data and assessing flexibility with 3D variability analysis (3DVA^30^)(Figure 1b, Supplementary Videos 1-3), cases where published crystal structures had multiple proteins in the asymmetric unit that could be assessed for consistency (Extended Data Figure 1c), and cases where only a cryoEM map was available (Extended Data Figure 1a,b).

Consistent with prior work^4^, triplicate 0.5 μs molecular dynamics simulations were effective at producing ensembles with reasonable correlation to what could be observed in the cryoEM data (Figure 1b, Extended Data Figure 1a,c), either demonstrating a similar range of motion to 3DVA or, for those fiducial markers that were too flexible for 3DVA (the megabodies 6 and 25^31^), predicting a substantial range of conformations. There does appear to be a consistent slight overestimation in the degree of motion, however this may be attributable to the MD simulations being performed at 27-30°C while cryoEM experiments typically begin at 4-25°C prior to flash freezing, which while rapid likely has some effect on conformational ensembles^32^. Despite the accuracy of the MD simulations, the computational expense highlighted a need for a more efficient protocol for design work.

We next evaluated how well AF2 conformer generation with multiple random seeds and MSA subsampling or enabling dropout at inference^19-21^ performed at predicting fiducial marker performance, based upon agreement with the cryoEM and/or MD simulation results. In general AF2-based sampling performed comparably to MD for recapitulating the protein dynamics seen in cryoEM (Figure 1b, Extended Data Figure 1a,b). This contrasts with other metrics, including pLDDT score in the region between fused domains, where we could identify examples of rigid constructs with poor pLDDT score (for example the frizzled receptor 5-BRIL fusion, Figure 1b) and flexible constructs with favorable pLDDT scores in linker regions (including Fab; where pLDDT score largely fails to predict the effect of adding disulfide crosslinks while the AF2-based ensemble captures the effect, Figure 1b). Analysis of the distribution of the number of structures with a given target-fiducial marker angle (Extended Data Figure 2a-d) suggests that most AF2-based approaches (including sampling multiple random number seeds without additional modification) qualitatively agreed with the 3DVA and/or MD simulation results, but overly limiting the MSA component risked producing unlikely if not completely misfolded structures (Extended Data Figure 2e). AF2 with dropout enabled was chosen for prospective design as a balanced solution, avoiding spurious structures and potential MSA issues for our engineered proteins while enhancing sampling, although several of the tested parameters likely would have been equally effective. The only case examined where the AF2-based ensembles failed was the SMO-PGS2 fusion^6^ (Extended Data Figure 2f), which largely seems to result from a failure to predict the correct structure at all.

### High-Throughput Determination of Inactive-State GPCRs with Bespoke Fiducials

To test the robustness of our approach prospectively, we designed 13-25 kDa fusion proteins for four GPCRs, MRGPRX2, MC2R, V1AR, and GIPR. We produced 3-5 constructs for each receptor that were predicted to have a well converged ensemble from AF2 dropout sampling (Figure 2) for testing in fluorescence size exclusion chromatography (FSEC). The most biochemically well-behaved design based on FSEC for each receptor was expressed, purified, and imaged, with every design imaged in cryoEM yielding at least a 3.4 Å map (Figure 2a, Extended Data Figure 3). 3DVA further resolved little heterogeneity in the connection between the receptor and fiducial (Extended Data Figure 4a).

**Figure 2:**
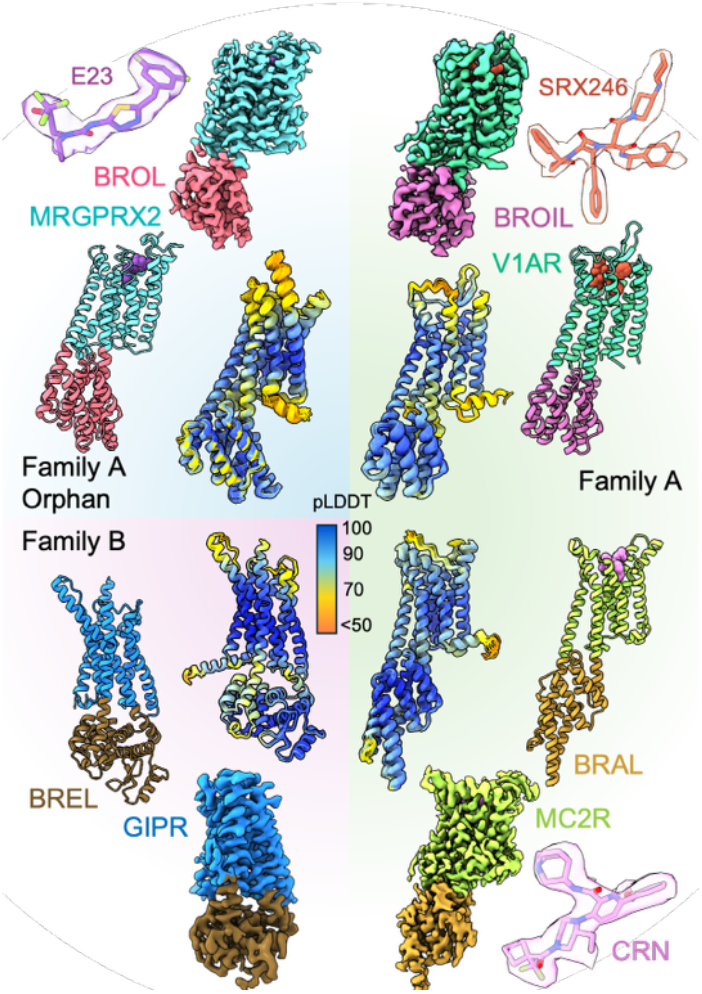
High-Throughput Determination of Inactive State GPCR Structures. CryoEM models and maps for the four receptors studied, with the fiducial marker colored separate from the receptor sequence. Map-model agreement is provided for small molecule antagonists. The ensemble of structures generated with AF2 with dropout used to assess rigidity is aligned and colored by pLDDT score (inner ring).

Each structure includes a bound antagonist (MRGPRX2-E23, V1AR-SRX246, a compound investigated for intermittent explosive disorder and PTSD, MC2R-CRN04894, an investigational oral compound for Cushing’s syndrome and congenital adrenal hyperplasia, and GIP-[N^α^-Ac, L14, R18, E21] hGIP_(5-31)_-K11 (γE-C16) denoted here as GIP(5-31*), a compound developed for *in vivo* pharmacology studies of weight loss), providing an opportunity to assess not only the structural pharmacology of these compounds but also the performance of recent AI-based ‘co-folding’ approaches for predicting the structure of protein-small molecule complexes. E23 binds to MRGPRX2 through interactions on transmembrane helix 3, 6, and 7 together with extensive hydrophobic interactions from ECL3-TM7, which forms a cap on top of E23 (Figure 3a). This involves a substantial rearrangement of TM7 at proline P7.36 compared to recently resolved active-state cryoEM structures with peptide and small-molecule agonists (PDB:7VV6, 7S8L^33,34^, Figure 3b,c), effectively converting the top of TM7 into an extension of ECL3. This also involves a rearrangement of the N-terminus and TM1 region that is obligate due to a disulfide bond between C7.32 and C26 (Figure 3b). While co-folding models generally place the ligand near the correct binding site with the correct orientation, the protein-ligand interactions are largely incorrect due to a failure to predict the change in fold at the top of TM7 (Figure 3c, Extended Data Figure 4b,c).

**Figure 3:**
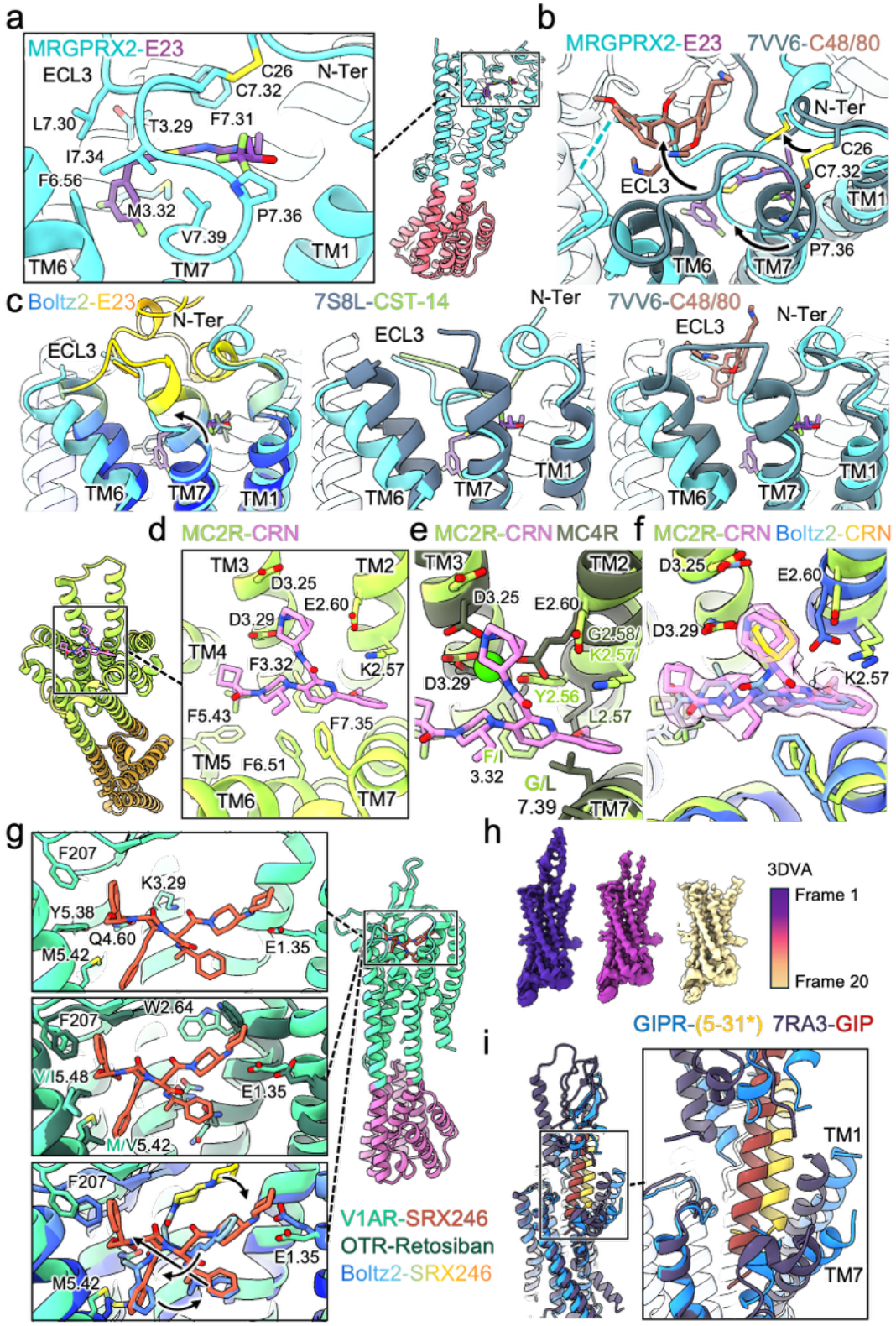
Receptor Conformational Changes Drive Antagonist Binding Yet are Poorly Predicted by Co-Folding Models. **a**, Inset of the MRGPRX2(aqua)-E23(purple) binding pocket key residues including those from ECL3-TM7. **b**, Comparison of our cryoEM structure with MRGPRX2 from PDB:7VV6 (green gray) bound to C48/80 (orange) with arrow depicting the rearrangement of TM7 to cap the antagonist binding pocket. **c**, Overlay of the ECL3 region of our MRGPRX2 structure with the top Boltz-2 predicted structure (left, colored by pLDDT), PDB:7SGL (center, blue gray) and PDB:7VV6 (right, green gray). **d**, Inset of the MC2R(lime)-CRN04894(pink) ligand binding site. **e**, Overlay of our cryoEM structure of MC2R with MC4R from PDB:6W25 comparing the position of the azaadamantane group with the calcium ion of MC4R (green sphere). **f**, Overlay of our cryoEM structure of MC2R and the CRN map with the top Boltz-2 predicted structure. **g**, Insets of our V1AR(mint)-SRX246(orange) structure (top), our structure overlaid with PDB:6TPK of OTR (dark green, middle), and our structure overlaid with the top Boltz-2 predicted complex with arrows depicting swapped moieties. **h**, Maps from 3DVA processing of GIPR. **i**, Overlay of GIPR-GIP(5-31*) (blue, yellow) and PDB:7RA3 of GIPR-GIP complex (dark Purple, brick).

CRN04894 binds to MC2R in what appears to be a cation-independent fashion, with the azaadamantane moiety forming a surrogate for occupying the canonical melanocortin calcium site with D3.25 and D3.29, while E2.60 interacts with K2.57 which forms a cation-π interaction with CRN04894 (Figure 3d). This contrasts heavily with the MC4R antagonist SHU9119, which was found in a crystal structure in complex with MC4R to require a calcium ion for binding like melanocortin receptor agonists (Figure 3e)^35^. Overlaying this structure with our MC2R structure reveals clear clashes with CRN04894 binding to MC4R due to residue substitutions (Figure 3e), including G7.39L and K2.57N (although this residue is L in PDB:6W25^35^). Further, K2.57, which coordinates both the ligand and E2.60, is geometrically more aligned with glycine G2.58 in MC4R, which likely contributes significantly to CRN04894 being a highly selective compound for MC2R. Co-folding approaches performed the best for CRN04894 binding to MC2R compared to the other small molecule cases examined, however a shift of the ligand in the pocket (perhaps due to TM2 being shifted in the TM bundle to more closely resemble previous melanocortin receptor structures, Figure 3f, Extended Data Figure 4b,c) results in a heavy-atom RMSD for the top ranked pose with Boltz-2 of ∼3.2 Å and a loss of the K2.57 cation-π interaction.

The binding pose of SRX246 is mostly driven by extensive hydrophobic packing across the V1AR pocket, with a salt bridge between one of the piperidine moieties and E1.35 (Figure 3g). The preferred binding of SRX246 for V1AR over OTR stems from a combination of substitutions such as M5.42V and V5.48I introducing steric clashes, a repositioning of TM2 and TM1 causing E1.35 to be unable to form a salt bridge with SRX246, and W2.64 shifting rotamers to block ligand binding (PDB:6TPK^36^, Figure 3g). Likely as a result of the largely hydrophobically-driven binding interface and seeming interchangeability of the interactions of some of the SRX246 moieties (e.g. numerous phenyl groups), co-folding methods generally predict the structure of V1AR fairly accurately yet perform the worst of all the ligands at predicting the bound pose of SRX246, exhibiting total rearrangement of the compound in the pocket (Figure 3g, Extended Data Figure 4b,c).

Our consensus refinement map for GIPR did not have well-resolved features for the peptide antagonist or extracellular domain (ECD)(Figure 2), although 3DVA and 2D class averages demonstrated the presence of particles both with and without an ordered ECD (Figure 3h, Extended Data Figure 4d). This suggests that the antagonist peptide GIP(5-31*) might primarily engage the ECD and only loosely interact with the TMD as a result of the significant N-terminal truncation. Performing classification using the ECD present and absent maps from 3DVA produced a particle stack which enabled a reconstruction with the peptide and ECD partially resolved. Comparing our structure of GIPR-GIP(5-31*) to the existing cryoEM structure of GIP-bound GIPR (PDB:7RA3^37^, Figure 3i) demonstrates that the antagonist peptide is shifted in the pocket by several angstrom, together with the tops of transmembrane helices 1, 6, and 7. The N-terminus of the antagonist peptide extends significantly less deeply into the binding pocket and is poorly ordered, consistent with a reduced affinity for the TMD (Figure 3i, Extended Data F.igure 4e). We were also able to resolve a map feature that likely corresponds to the lipidated lysine, which extends between TM1 and TM2 (Extended Data Figure 4f).

### An Extracellular Fiducial for β2AR Reveals Strong Agonists Almost Completely Activate Key Motifs for Transducer Binding

Next we selected the β2 adrenergic receptor (β2AR) as our system for generating an extracellular fiducial marker given its status as one of the most prototypical and well-studied family A GPCRs with numerous well characterized ligands and extensive exploration with spectroscopic studies^7-11^. Given the challenges in designing a TM1 fusion and a desire to assess how much flexibility would be tolerated, five designs were selected spanning total predicted rigidity to a modest degree of predicted conformational heterogeneity. Based on FSEC we identified two designs, denoted β2BB3 and β2BB5, that were particularly well behaved biochemically (Extended Data Figure 5a), and testing in TRUPATH BRET assays demonstrated signaling similar to the wildtype receptor, with an EC50 for epinephrine within one log unit of the WT β2AR for both BB3 and BB5 (Extended Data Figure 5b). β2BB3 was predicted to be one of the most well converged ensembles while β2BB5 was predicted to be the most flexible design selected (Figure 4a), and consistent with the predictions 2D class averages from small screening datasets performed on a 200 kV microscope suggested the fiducial for β2BB3 to be sufficiently rigid for alignment while β2BB5 was not (Figure 4b).

**Figure 4:**
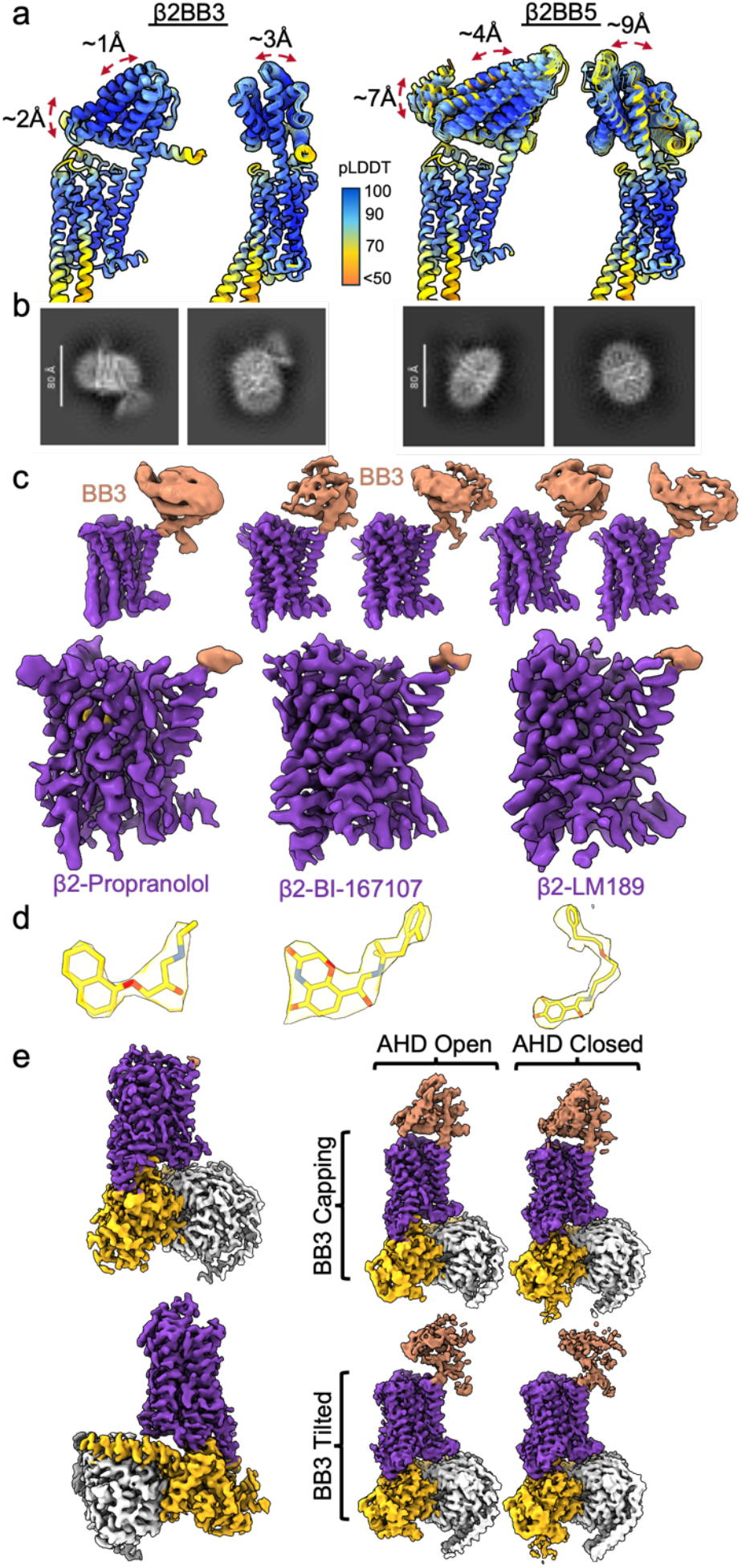
An Extracellular Fiducial Marker Enables Ligand-Agnostic Structure Determination of a Family A GPCR. **a**, AF2 dropout ensembles of β2BB3 and β2BB5 aligned on the β2 transmembrane domain and colored by pLDDT score. **b**, 2D class averages of β2BB3 and β2BB5 from ∼800 screening images collected on a 200 kV microscope. **c**, CryoEM maps of β2BB3 (purple receptor, beige fiducial) obtained with propranolol, BI-167107, and LM189 with unsharpened maps derived from 3DVA depicted on top and sharpened local refinement on the receptor on bottom. **d**, CryoEM maps for the three ligands from the local refinement maps. **e**, CryoEM map of the β2BB3-BI-167107-Gs complex together with four unsharpened maps depicting the four primary conformational states that could be resolved with 3D classification (alpha helical domain open versus closed, BB3 capping vs tilted).

The β2BB3 construct enabled us to obtain cryoEM reconstructions bound to the prototypical antagonist propranolol, the extremely potent and efficacious agonist BI-167107, the Gi-based agonist LM-189^11^, and the complex of β2BB3-BI-167107 with wildtype Gs protein (Figure 4c,d,e). While sufficiently rigid for determining structures of the receptor TM bundle alone up to 3.1 Å resolution, the BB3 fiducial occupies two different conformations, one which matches the design model while the other is tilted by ∼40°, a geometry that may only be allowable due to the detergent micelle (Figure 4c,e). No structure prediction algorithm tested was able to produce this conformation (Extended Data Figure 5c), and MD simulations started from the tilted conformation in a lipid bilayer rapidly converted to the predicted conformation (Extended Data Figure 5d), further suggesting the detergent environment plays a role in stabilizing the alternative state. The dataset of β2BB3-Gs complex possessed further conformational heterogeneity in the Gs protein with both alpha helical domain (AHD) open and closed nucleotide-free states present (Figure 4e), consistent with previously reported work for β2AR-Gs^38^.

Comparing the structures of β2BB3 bound to propranolol, BI-167107, and BI-167107 in complex with Gs reveals that the high efficacy agonist BI-167107 induces almost complete outward movement of TM6 even in the absence of G-protein, with the introduction of the Gs α5 helix shifting TM6 by less than 1 Å further outward at most positions (Figure 5a). This contrasts with interpretations of prior double electron-electron resonance (DEER) work, which have suggested TM6 is shifted by several angstroms further upon complexation with Gs^9,11^. Based on our structures, the discrepancy likely arises from the positioning of the labels used in DEER (and single molecule Förster resonance energy transfer (smFRET), which also suggests further motion of TM6 upon G-protein binding), which are typically placed on C6.27 or L6.28C; both of which would occur almost one helical turn lower than the extent of TM6 that is well resolved in our agonist-bound cryoEM maps (Extended Data Figure 5e). The DEER and smFRET results then represent the convolution of TM6 movement and the flexibility of the end of ICL3. To probe this hypothesis, we performed MD simulations of BI-167107-bound β2AR with the TM5-ICL3-TM6 region modeled either similar to our structure with ICL3 added in a disordered loop, or with a highly extended TM6 helix (Figure 5b). In those simulations where TM6 was extended in a helix, the TM4-TM6 distance distribution was tightly peaked at ∼39 Å, corresponding to an open TM6 state (Figure 5b). Simulations starting from a structure based on the cryoEM result possessed a slightly shifted peak at ∼38 Å with a substantial shoulder at lower values corresponding to ICL3 bending back towards the intracellular cavity (Figure 5b), which more closely matches previously reported DEER results^11^. Introduction of Gs (particularly the α5 helix) to β2AR bound to a highly efficacious agonist thus displaces ICL3 out from under the receptor and/or induces additional folding of TM6, rather than inducing an outward movement of TM6 (Figure 5a), with ICL3 likely acting as an allosteric modulator of transducer coupling (consistent with what has been proposed based on FRET studies of ICL3 motion for β2AR^39^). This is further supported by 3DVA, which for the β2BB3-BI-167107 complex reveals that the primary heterogeneity in TM6 is in the degree of extension of the helix rather than inward and outward movement (Figure 5c). In some frames the map barely extends to include H6.31, while this region is well resolved through H6.31 in the β2BB3-BI-167107-Gs complex, at which point TM6 also appears to terminate (Extended Data Figure 5f,g). This also contrasts with the inactive state bound to propranolol, where TM6 appears to be slightly more extended, although the unsharpened map possesses a feature that may correspond to ICL3 threading backwards towards ICL2 (Extended Data Figure 5h).

**Figure 5:**
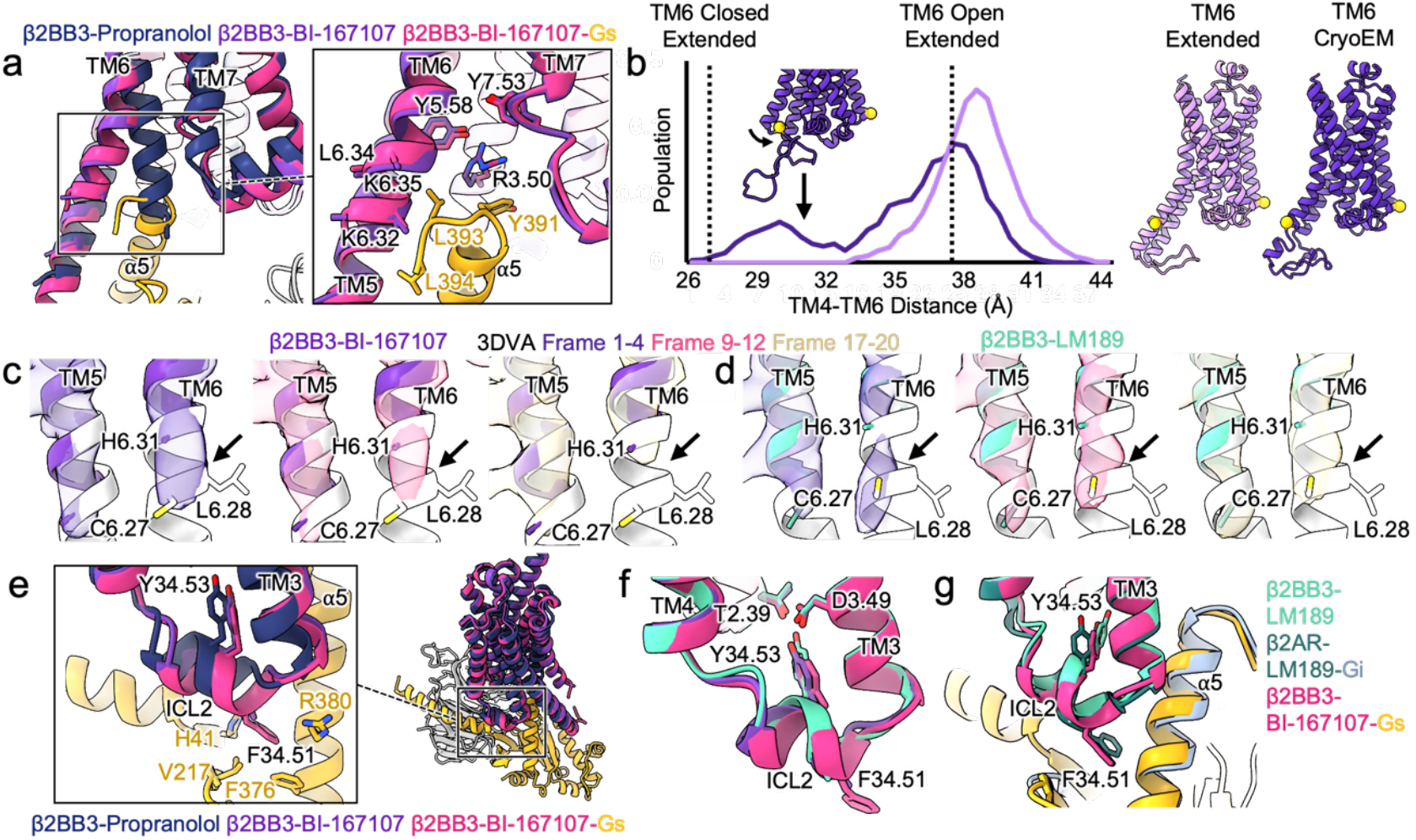
Strong Agonists Pre-Organize the β2 Intracellular Vestibule For G-Protein Coupling. **a**, Alignment of β2BB3 bound to propranolol (navy), BI-167107 (purple), and BI-167107 in complex with Gs (magenta receptor, goldenrod G-protein) with inset highlighting TM6 and key activation residues in the BI-167107-bound receptor with and without Gs. **b**, MD simulation results for the distribution of TM4-TM6 distances measured between N4.40 and L6.28 backbone nitrogens for β2BB3-BI-167107 starting from a model with an extended transmembrane helix 6 (light purple) or matching what is resolved in the cryoEM structure (purple). Dashed lines indicate TM4-TM6 distances corresponding to a fully open and fully closed extended TM6. **c**,**d**, 3DVA-derived maps demonstrating the heterogeneity in TM6 extension observed with β2BB3-BI-167107 (**c**) and β2BB3-LM189 (**d**). CryoEM structures (purple BI-167107, teal LM189) are overlaid with an idealized extended TM6 structure (white). **e**, Comparison of ICL2 in the β2BB3 propranolol, BI-167107, and BI-167107 Gs complex. **f**, Comparison of ICL2 for β2BB3 bound to BI-167107 (purple), BI-167107 with Gs (magenta), and LM189 (teal). **g**, Overlay of ICL2 from β2BB3-LM189 (teal), β2AR-LM189-Gi complex (dark green receptor, slate Gi), and β2AR-BI-167107-Gs (magenta receptor and goldenrod Gs).

In contrast to prior crystal structures of β2AR, ICL2 forms a helical structure in the inactive state bound to propranolol (Extended Data Figure 6a), raising the possibility that the previously observed loop is an artifact of crystal packing interactions (Extended Data Figure 6b), although it is possible the loop may represent a higher energy alternative conformation. Comparing the ICL2 region (one of the key motifs for binding G-protein) of the β2BB3-propranolol structure to β2BB3-BI-167107 with and without Gs reveals that BI-167107 alone is again sufficient to induce a substantial shift in the motif to nearly match the Gs-bound result (Figure 5e). To assess whether there were any structural features that might indicate how LM189 achieves Gi bias, we aligned the β2BB3-BI-167107 structure to that with β2BB3-LM189, which reveals a largely superimposable structure for much of the receptor (Extended Data Figure 6c). However, a subtle shift in ICL2 could be modelled (to within the resolutions obtained, Figure 5f, Extended Data 6d), consistent with prior work indicating this motif likely plays a role in the greater propensity of LM189 to signal through Gi^11^. Superimposing our structures with a published LM189-bound β2AR-Gi complex structure (PDB:9BUY^11^) demonstrates that ICL2 in the Gi complex is in a unique position from any of our resolved structures including β2BB3-LM189 (Figure 5g), although it is also influenced by a positive allosteric modulator that may further affect ICL2 conformation. While consensus maps for BI-167107 and LM189 were similar for TM6, 3DVA suggested TM6 may have a slightly greater propensity to be extended further with LM189 (Figure 5d), which is also the case in the LM189-Gi complex (Extended Data Figure 6e) and may also contribute to the bias of LM189. This would also be consistent with DEER observations suggesting longer TM4-TM6 distances with LM189 than BI-167107^11^ as the tagged site is more likely to be in a helical region. While co-folding approaches were able to reliably model the ligand bound pose for propranolol and BI-167107, some models struggled with the terminal phenyl ring of LM189 (Chai-1, OpenFold 3) (Extended Data Figure 7a). Further, TM6 was predicted to either occupy an entirely inward conformation (Boltz-2, OpenFold 3) or outward (Chai-1) regardless of ligand identity (Extended Data Figure 7a), failing to capture the core activation responses observed here.

## Discussion

Here, we demonstrate that combining generative protein design with a machine-learning based protein structure ensemble prediction method enables reliable production of GPCR fiducial markers, even when fused to traditionally intractable sites such as TM1. This strategy allowed high-throughput determination of inactive state structures of several pharmacologically important receptors and established a platform for imaging β2AR with any arbitrary ligand while leaving the intracellular vestibule of the receptor completely native. These β2AR structures suggest that, in contrast to some existing models of GPCR activation, the receptor in the presence of a sufficiently potent agonist undergoes almost complete rearrangement of the ordered domains to match the G-protein bound form. In this framework the heterogeneity resolved in spectroscopic studies may arise largely from motions in the dynamic intracellular loop elements and how they relate to the surrounding ordered domains. This interpretation is consistent with emerging views that aspects of precise pharmacology (partial versus full agonism, biased agonism) involve subtly influencing these dynamic intracellular elements, which in turn regulate transducer coupling^39,40^. We note, however, that these cryoEM structures may capture subsets of lower energy state(s) with a given ligand and higher energy structures may exist at room temperature, especially given that cryoEM vitrification is not instantaneous.

All of the fiducial designs presented here should be transferable to other receptors, which can be easily tested computationally with our AF2 dropout approach. Our extracellular fiducial marker can also likely be grafted to other receptors, although due to the substantial structural variety of GPCR extracellular pockets further designs will likely be needed in many cases. We anticipate pharmacologically inert extracellular fiducial markers for GPCRs will aid structural pharmacology and structure based drug discovery efforts and further facilitate obtaining structural snapshots of the dynamic process of GPCR signaling. Enabling reconstructions of the receptor alone will allow for improved tracking and visualization in several contexts, including studying the association of GPCRs with their signaling partners in a time-resolved fashion (as has been done with looking at the GTP-induced G-protein activation and dissociation of formed complex^38,41^). This will also benefit attempts to visualize proposed tertiary and larger complexes of GPCR, signaling partners, and additional receptors and/or downstream signaling elements (referred to as ‘signalosomes’ or ‘HOTS’^42,43^).

Finally, although we applied this framework exclusively to GPCRs, the approach should facilitate cryoEM structure determination for a broad range of proteins. By avoiding the need to screen for antibodies or nanobodies and enabling constructs to be tested computationally prior to cryoEM data collection, this pipeline will help streamline structural interrogation and structure-based drug discovery of small, dynamic targets. Obtaining snapshots of these systems remains incredibly valuable, as this present work has highlighted that traditionally difficult structural biology targets including GPCRs remain challenging for co-folding models, which struggle to accurately predict both ligand binding poses and ligand-induced conformational changes. Improving our ability to access these states will provide the crucial training data for the next generation of predictive models for co-folding and ensemble generation, a rapidly improving space^44,45^, where more accurate methods could replace the AF2 dropout component of our approach to even further increase reliability.

## Supporting information

All Supplemental Figures

## Acknowledgements

NIH R00 HD107581 (M.J.R.), CPRIT award RR230042 (M.J.R.), M.J.R. is a CPRIT Scholar in Cancer Research. We thank Bharti Singal for cryoEM data collection at the Stanford University Cryo-EM center (cEMc). We thank Dr. Gaya P. Yadav for cryoEM data collection at the Laboratory for Biomolecular Structure and Dynamics (LBSD) of Texas A&M University. The LBSD is supported, in part, by the Department of Biochemistry & Biophysics, AgriLife, and the Texas A&M University. We thank Mehdi Zarei for cryoEM data collection at the Electron Cryo-Microscopy Core Facility at UTHealth (NIH S10ODO32204). We thank Dr. Erik Hartwick for cryoEM data collection at the Biochemistry Krios Electron Microscope Facility (BioKEM) at CU Boulder (RRID:SCR_019057).

## Contributions

A.V. purified proteins and cloned constructs. M.J.R. conceived and supervised the project, designed proteins, expressed and purified proteins, prepared cryo-EM grids, collected, analyzed and processed cryo-EM data, built and refined atomic models, designed and executed molecular dynamics simulations, analyzed data, and prepared figures. Y. S. performed BRET assays and surface expression testing under the supervision of J.G.E. M.J.R. wrote the manuscript with input from J.G.E..

## Conflict of Interest

The authors have no conflict of interest to declare.

## Data availability

The atomic coordinates have been deposited in the Protein Data Bank Cryogenic electron microscopy maps have been deposited in EMDB

## Methods

### CryoEM Dataset Reprocessing

Datasets for the fiducial marker-target complexes/fusions were pulled down from EMPIAR with accession codes FZD5 EMPIAR-10507^5^, Fab EMPIAR-10380^45^, 6DS-Fab EMPIAR-12290^28^, Mb38 EMPIAR-10912^47^, Mb6 EMPIAR-11135^4^. Data was imported into cryoSPARC^48^ where it was processed with patch motion correction, patch CTF estimation, a single round of 2D classification followed by iterative rounds of *ab initio* reconstruction with multiple classes and heterogeneous refinement with those maps until all junk particles were removed. Non-uniform refinement followed by local refinement on the static component targeted by the fiducial marker was subjected to 3DVA^30^ of the target-fiducial system to assess fiducial flexibility.

### Molecular Dynamics Simulations

For MD simulations to assess flexibility of published fiducial markers, systems were prepared from deposited PDB structures when a full structure was available (FZD5, Fabs, megabodies with crystal structures, macrobodies with crystal structures) or AF2 predictions when it was not (some megabodies). Simulations of β2AR/β2BB3 were initiated from our cryoEM structures. Systems were prepared with CHARMM-GUI^49^ solubilized in TIP3P water^50^ with a 95% POPC/5%CHS lipid bilayer oriented with the PPM webserver^51^ when the protein was a membrane protein. Systems were generally simulated in 150 mM NaCl, with the exception of active-state GPCRs which were simulated in 100 mM NaCl. GPCRs were also lipidated on helix 8 when lipidation site(s) were present. Simulations were prepared with the CHARMM36 forcefield^52-54^, for simulation in NAMD^55^. Molecular dynamics was performed with a Langevin thermostat and Nose-Hoover Langevin piston barostat at 1 atm with a period of 150 fs and decay of 75 fs and periodic boundary conditions with nonbonded interaction smoothing at 10 Å to 12 Å with long-range interactions handled with particle mesh Ewald. A 2 fs timestep was employed with SHAKE and SETTLE algorithms used. All non-hydrogen, non-water/ion atoms were restrained with harmonic restraints of 1 kcal/mol/Å^2^ and the systems were minimized for 1,500 steps before gradual heating from 0 to 303.15K in 20K increments with 0.4 ns of simulation per increment, with 1 kcal/mol/Å^2^ harmonic restraints applied to all non-hydrogen, non-water/ion atoms for membrane simulations. Systems without a membrane were simulated for 45 ns as equilibration before 500 ns of simulation as production. For systems with a membrane, 10 ns of equilibration with 1 kcal/mol/Å^2^ restraints on all protein heavy atoms was followed by 10 ns of equilibration with 1 kcal/mol/Å^2^ restraints on all protein CA atoms with subsequent rounds of 10 ns of equilibration with 0.5 and 0.3 kcal/mol/Å^2^ CA restraints, followed by an additional 20 ns of unrestrained equilibration before production. 500 ns of production was employed for fiducial marker simulations while 1 μs of simulation for the β2AR/β2BB3 simulations of TM6 movement. All β2AR/β2BB3 simulations were performed with 5 replicates while all other simulations had 3 replicates. TM4-TM6 distance was measured with VMD^56^ taking the distance between the backbone nitrogen of L6.28 and N4.40.

### AF2-Based Ensemble Generation and Protein Design

AF2^15^ implemented in ColabFold^20^ was used to evaluate different approaches for ensemble generation with MSA modulation. For extensive sampling of fiducial marker-linker-target angle (Extended Data Figure 2), 50 random number seeds were used for each sampling approach, which included default AF2, the use of dropout, and limiting the depth of the MSA sampling ranging from 16:32 to 256:512. For all other uses 3 random number seeds were employed. Fusion proteins were designed in RFdiffusion^13^. Designs were typically generated in 2-5 rounds of 50-150 residues. Sidechain design was performed with ProteinMPNN^23^ with the sequence for the receptor held fixed. Ensemble prediction for the designed sequences was performed with AF2^15^ implemented in ColabFold^20^ with dropout enabled and three random number seeds (approximately 5-10 minutes per design on a single GPU). The predicted ensemble of structures was aligned and assessed for a low RMSD, particularly amongst the top scoring ∼10/15 models although strong preference was given to designs where all 15 models were tightly converged. For GIPR the same procedure was performed with a marginally relaxed criteria for convergence (due to the predictions producing almost exclusively active conformations of TM6) followed by AlphaFold-Multistate^58^ prediction with 5 random seeds and an inactive-GPCR bias to generate an ensemble for an inactive TM6 conformation with more stringent acceptance.

### Expression and Purification of GPCR Fusions

Pellets containing receptor-fiducial marker fusions cloned into standard expression vectors (N-terminal hemagglutinin, FLAG epitope, β2AR N-terminus, TEV cleavage site; C-terminal C3 cleavage site, GFP, 6x His tag) were expressed in Sf9 cells, harvested via centrifugation, and snap-frozen in liquid nitrogen for storage (The exception being the β2 constructs, which were cloned with the native N-terminus and no cleavage site, with the fiducial marker if present inserted in the N-terminal tail at the designed location). To start purification, the pellets were thawed and lysed with a hypotonic lysis buffer containing 10 mM HEPES pH 7.5, Pierce Universal Nuclease, protease inhibitor cocktail, 1 mM MgCl_2_, 100 µM TCEP, 1 mM EDTA, and 1 mM benzamidine, and stirring was allowed at 4°C for 1 hour. Lysis buffer and all subsequent buffers contained 10 µM antagonist/agonist for the given receptor and preparation. Membranes were harvested by ultracentrifugation at 100,000xg for 35 minutes, and pellets containing the membranes were resuspended in solubilization buffer containing 20 mM HEPES pH 7.5, 500 mM NaCl, 1 Mm MgCl_2_, Pierce Universal Nuclease, 100 µM TCEP, 1 mM benzamidine, and protease inhibitor cocktail. Detergent was added dropwise at 4°C while stirring, to a final concentration of 1% LMNG/0.1% CHS/0.1% Sodium Cholate. Following 3 hours of gentle stirring, ultracentrifugation at 100,000xg was done to remove insoluble debris. Solubilized receptor was supplemented with 20 mM imidazole and loaded over a nickel resin column that had been washed with 10 column volumes of buffer containing 20 mM HEPES pH 7.5, 500 mM NaCl, 20 mM imidazole, and 0.1% LMNG/0.01% CHS. The column was washed with the same wash buffer and the protein was eluted from the column with 2 column volumes of buffer containing 20 mM HEPES pH 7.5, 250 mM NaCl, 10% glycerol, 250 mM imidazole, and 0.1% LMNG/0.01% CHS. Eluted protein was supplemented with 5 mM CaCl_2_ and loaded over a M1 flag resin column that has been washed with 3 column volumes of buffer containing 20 mM HEPES pH 7.5, 250 mM NaCl, 2 mM CaCl_2_, and 0.01% LMNG/0.001% CHS. The column was washed with the same wash buffer and purified receptor was eluted with 2 column volumes of buffer with 20 mM HEPES pH 7.5, 1 mM EDTA, 250 mM NaCl, 0.01% LMNG/0.001% CHS, 10% glycerol, and flag peptide. Following elution, the purified receptor was supplemented with 100 µM TCEP and treated with HRV-3C protease overnight before concentration in a 50 kDa molecular weight cutoff spin concentrator, and subsequently subjected to size-exclusion chromatography with a Superdex 200 column in a buffer containing 20 mM HEPES pH 7.5, 150 mM NaCl (inactive state) or 100 mM (active state), and 0.001% LMNG/0.0001% CHS/0.00033% GDN, 100 µM TCEP and 100 µM antagonist/agonist. The preparation of GIPR was identical with the exception of the removal of TCEP, 1 µM ligand utilized for preparatory steps, and the C-terminal GFP/tag was not cleaved.

### Expression and Purification of Gs Heterotrimer

Expression of Gs heterotrimer was performed as previously described^57^. Purification of Gs heterotrimer began with thawing and resuspending cell pellets in hypotonic lysis buffer containing 20 mM HEPES pH 7.5, 1 mM MgCl_2_, 5 mM β-Mercaptoethanol, 100 µM GDP, 1 mM EDTA, 5% glycerol, Pierce Universal Nuclease, and protease inhibitor cocktail, followed by gentle stirring for 30 minutes at 4°C. Lysate was then spun down at 100,000xg for 35 minutes, and pellets were resuspended in a solubilization buffer containing 20 mM HEPES pH 7.5, 5% glycerol, 1 mM MgCl_2_, 5 mM β-Mercaptoethanol, 100 mM NaCl, 1% sodium cholate hydrate, 100 µM GDP, Pierce Universal Nuclease, and protease inhibitor cocktail, and allowed to stir at 4°C for 1 hour. A second round of ultracentrifugation proceeded at 100,000xg for 35 minutes. Solubilized protein was supplemented with 30 mM imidazole and incubated with Ni-NTA beads for 1 hour on ice. Beads were harvested by centrifugation at 300xg, packed into a column, and washed with a series of buffers containing 30 mM imidazole and 50% solubilization/50% E2 buffer, 25% solubilization/75% E2 buffer, 12.5% solubilization/87.5 E2 buffer, and 100% E2 buffer (E2 buffer contained 20 mM HEPES pH 7.5, 1 mM MgCl_2_, 100 mM NaCl, 5% glycerol, 100 µM GDP, 5 mM β-Mercaptoethanol, and 0.05% LMNG/0.005% CHS). The protein was eluted from the column with 3 column volumes of buffer containing E2 buffer and 250 mM imidazole, and was subject to incubation with 1 mg of 3C protease per 50 mg of protein along with overnight dialysis at 4°C in a 3,500 MWCO dialysis tubing. The following day, the column was washed with 10 column volumes of buffer containing E2 and 30 mM imidazole, and the overnight cleavage was loaded over the nickel, and flow through was collected. An additional 2 column volumes of E2 buffer and 30 mM imidazole was loaded over the column, and flow through was collected. Eluted protein was concentrated to 500 µL in a 30 kDa MWCO spin concentrator prior to being subject to SEC chromatography with a Superdex 200 column in buffer containing 20 mM HEPES pH 7.5, 1 mM MgCl_2_, 20 µM GDP, 100 µM TCEP, 100 mM NaCl, 5% glycerol, and 0.01% LMNG/0.001% CHS.

### Complex Preparation

β2BB3-BI-167107-Gs complex was prepared by incubating purified β2BB3-BI-167107 with Gs for 1 hour on ice, followed by addition of apyrase and incubation overnight. Successful complex was pulled down with an M1 flag resin column with an equilibration/wash buffer containing 20 mM HEPES pH 7.5, 100 mM NaCl, 2 mM CaCl_2_, 10 μM BI-167107 and 0.005% LMNG/0.0005% CHS. Purified receptor was eluted with 2 column volumes of buffer with 20 mM HEPES pH 7.5, 1 mM EDTA, 100 mM NaCl, 0.005% LMNG/0.0005% CHS, 10% glycerol, 10 μM BI-167107 and flag peptide. Complex was supplemented with 100 µM TCEP, concentrated, and subjected to size exclusion chromatography on a Superdex 200 column into a final buffer of 20 mM HEPES pH 7.5, 100 mM NaCl, 0.001% LMNG/0.0001% CHS/0.00033% GDN, 100 µM TCEP and 100 µM BI-167107.

### CryoEM Sample Preparation & Data Collection

Samples were concentrated to 5-10 mg/ml for cryoEM grid preparation. R1.2/1.3 UltrAufoil 300 mesh grids were glow discharged in a Pelco unit for 45 seconds at 10 mA. Grids were plunge frozen in a Vitrobot Mark IV after addition of 3 µL of sample. ICL3 fusion data was collected on a G3 Titan Krios equipped with a Falcon4i/SelectrisX energy filter. ICL2 fusion data was collected on a G3 Titan Krios equipped with a Falcon 4i detector and Selectris energy filter. β2BB3 data was collected on a G4 Titan Krios equipped with a K3 direct electron detector and a Bioquantum energy filter. Glacios screening data was collected on a Glacios 1 equipped with a Falcon 4i detector and Selectris X energy filter. Details of each data collection can be found in Extended Data Tables 1 and 2 and Supplementary Table 1.

### CryoEM Data Processing

All data was processed in cryoSPARC^48^. Raw movies were aligned with patch motion correction and CTF estimation was performed with patch CTF estimation. Particle picking was performed with template-based picking using templates for GPCR-G-protein complexes or GPCR with an ICL3 fiducial marker. Extracted particles were subjected to 2D classification to remove noise particles. Further cleaning was performed with iterative rounds of *ab initio* model generation with multiple classes and heterogeneous refinement. Once particle stacks were largely free of junk particles, non-uniform refinement was applied to obtain reconstructions. Several reconstructions were found to produce a strong appearance of preferred orientation from non-uniform refinement (based on 3D FSC and visual inspection of the map) despite a balanced appearing Euler angle distribution and absence of these features in previous processing steps. This was improved with the use of the HR-HAIR approach^59^, which allowed for the removal of further subtle inferior particles and a high-resolution initial model without orientational artifacts. 3DVA was run on final reconstructions to probe for further heterogeneity, and the first principal component analyzed. Local refinement on the transmembrane region was performed in several cases to further improve the GPCR/ligand resolution. In the case of GIPR, final heterogeneous refinement was performed with a class with and without the ECD present (obtained from 3DVA), which resulted in a final reconstruction with some resolution of the bound antagonist peptide and ECD.

### CryoEM Model Building

The AF2 predicted structure was docked for all initial structures of fiducial marker complexes, while PDB:8DGZ^38^ was also used for the Gs complex. Manual model building was performed in Coot^60^ with refinement in Phenix^61^. Initial ligand placement was performed with GlideEM^62^. Details of cryo-EM map and model refinement are found in Extended Data Tables 1 and 2. Euler angle distribution plots, FSC curves, local resolution plots, and selected map-model agreement panels are provided in Extended Data Figures 1 and 2 and Supplementary Figures 1 through 4.

### BRET Assays

HEK293 cells were seeded at a density of 2 × 10^6^ in 10mL DMEM in 10-cm plates overnight. Cells were transiently co-transfected with 1480ng of TRUPAHT triple sensor and 296ng receptor (5:1 ratio) using TransIT-2020 (Cat#MIR 5406, Mirus bio) following the instructions and incubated at 37°C, 5% CO2, >90% humidity overnight. After incubation, cells were harvested and resuspended in 1% dialyzed FBS DMEM. Cells were then plated at a density of 20,000 cells/40μL per well in the PDK (Sigma, P6407-5MG)-coated, clear-bottom 384-well assay plates (Greiner Bio-One, 781098) overnight. Assay buffer (20 mM HEPES in 1x HBSS, pH to 7.4 with KOH) and 3x drug buffer (assay buffer + 3mg/ml BSA, 0.3 mg/ml ascorbic acid) were prepared at least 2 hours before the experiment. rugs were diluted to 3x working concentration in 3x drug buffer using 96-well plates (Fisher, 12-566-120). Drug concentrations were created by serial dilution at half-log concentrations. The bioluminescent substrate Prolume purple (cat #369, NanoLight) was prepared in nanofuel (cat #399, NanoLight) to a concentration of 1 mM and stored at −80 °C. The growth media was flicked from each well, and a white backing (PerkinElmer, 6005199) was applied to the plate bottom. A concentration of 7.5 uM of the bioluminescent substrates, Prolume purple, was diluted in assay buffer and each well of the plates was added 20uL of the substrate. Subsequently, 10 μL of the drug solution was added to the well, producing a final well concentration of 5 μM. he plate could be read by the PheraStar Reader. The optic settings on the PheraStar Reader were set to have one multichromatic and simultaneous dual emission. The well scan was set to have a 3mm spiral average. The top optic was selected in the Optic window. The setting time was 0.1s in the general settings. In the kinetic window, four cycles were read; the start time was 0s, and the interval time was 0.5s. All the wells were selected to be read, no matter how many were occupied. One cycle to read the entire plate was 335s. The time to normalize results was set to 0, and the pause before the next cycle was 1 second. In the measurement panel, the gain settings were set to 3800 for Gain A and 3000 for Gain B channels. Gain adjustment has a value of 40% for both the A and B channels. Cycle two data were used for analysis, and concentration–response curves were fit to a three-parameter logistic equation using GraphPad Prism.

### ELISA

Cell surface expression of β2AR wildtype, BB3, and BB5 was confirmed using immunohistochemistry. Cells were plated on 384-well plates at a density of 10,000 cells per well. Fixation was performed with 20 μl per well of 4% paraformaldehyde (Fisher, #AAJ19943K2) for 10 minutes at room temperature. Following fixation, cells were washed twice with 40 μl per well PBS. Blocking was carried out with 20 μl per well of 5% normal goat serum (Vector Laboratories, #S-1000) in PBS for 30 minutes at room temperature. After blocking, 20 μl per well of monoclonal ANTI-FLAG M2-Peroxidase (HRP) antibody (Sigma-Aldrich, A8592), diluted 1:10,000 in PBS, was added and incubated for 1 hour at room temperature. This step was followed by two washes with 80 μl per well PBS. Subsequently, 20 μl per well of SuperSignal ELISA Pico Chemiluminescent Substrate (Sigma-Aldrich, #37069) was added and incubated at room temperature for 5 minutes, and luminescence was measured using a PheraStar reader. Data were plotted as luminescent units (RLU).

## Notes

### Competing Interest Statement

The authors have declared no competing interest.

### Summary of Updates

The results section has been revised to include an additional reference.

