## Supplementary material for "Designing Rigid Protein Fiducials to Visualize GPCR Conformational States": All Supplemental Figures

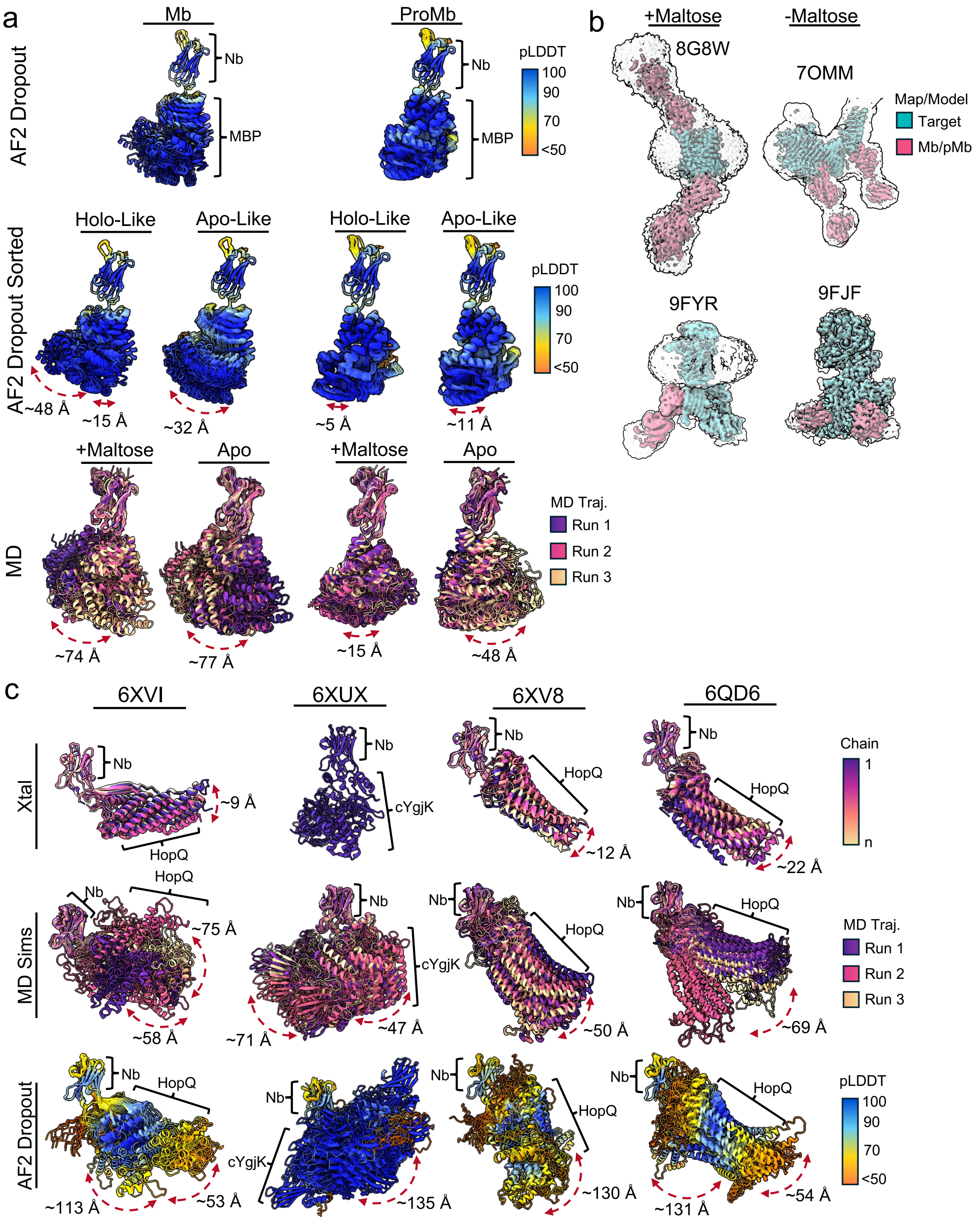


**Extended Data Figure 1: Fiducial marker systems with no deposited cryoEM micrographs a)** AF2 dropout ensembles for Macrobody and Pro-Macrobody joint (top) or split by apo and holo MBP conformations (middle) and equally spaced snapshots from triplicate MD simulation ensembles with and without maltose (bottom), highlighting only the Pro-Macrobody bound to maltose (holo) is predicted to be particularly rigid. **b)** CryoEM maps of Macrobody and Pro-Macrobody complexes at two contours (tight in color and loose in transparent gray) grouped by presence or absence of maltose. Consistent with the predictions, maltose-bound Macrobodies are more rigid. **c)** Megabody designs with deposited crystal structures; multiple chains in the asymmetric unit are aligned on the nanobody portion. Results are compared with MD simulations and AF2 dropout.

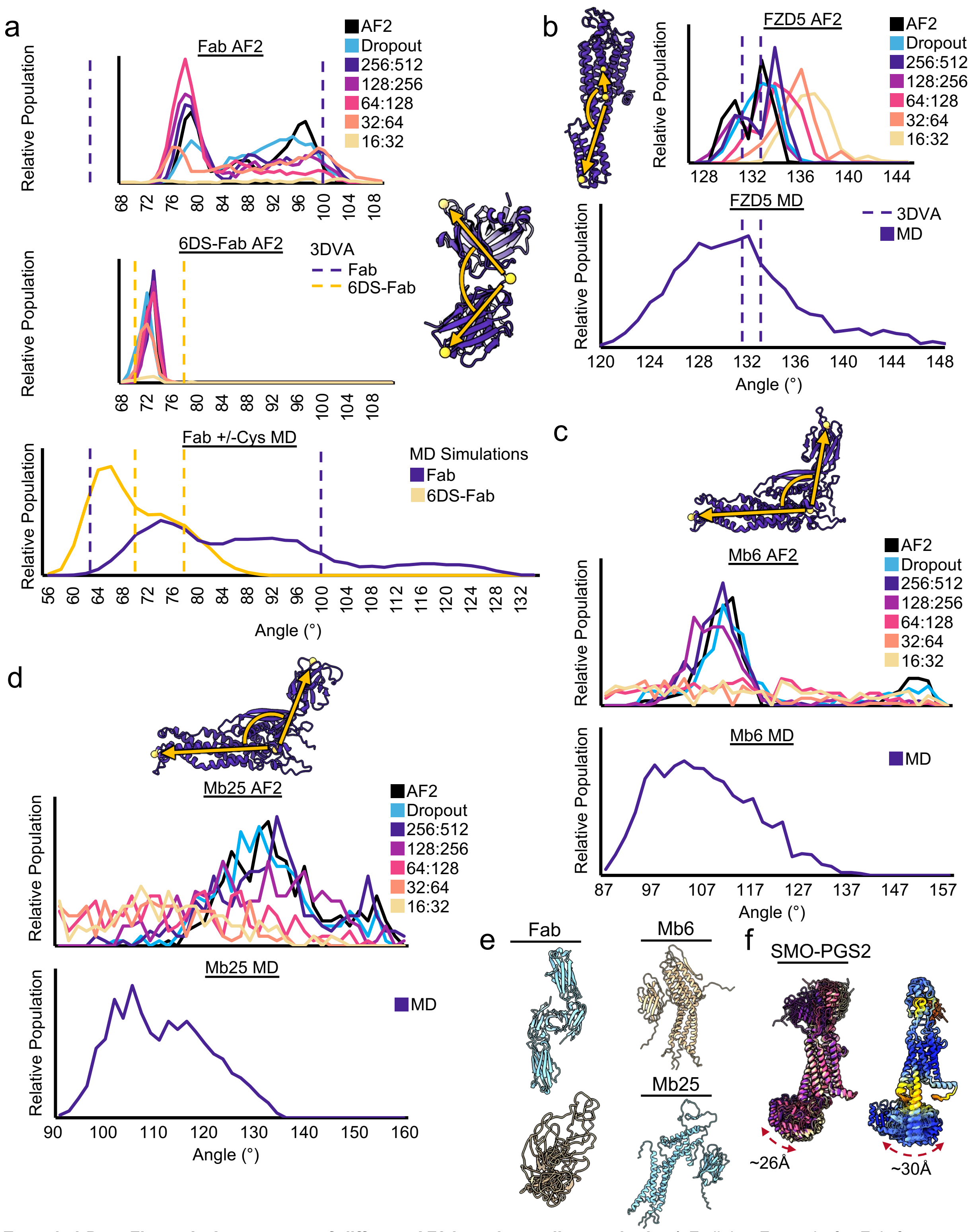


**Extended Data Figure 2: Assessment of different AF2-based sampling methods a)** Fv-linker-Fc angle for Fab fragment with and without cysteine rigidification from AF2-based sampling and MD. Dashed lines indicate the extrema observed from 3DVA of cryoEM datasets, misfolded structures were omitted from analysis. **b)** FZD5-linker-BRIL angle from AF2-based sampling and MD. Dashed lines indicate the extrema observed from 3DVA of the cryoEM dataset. Misfolded structures were omitted from analysis. **c)** Mb6 Nb-linker-HopQ angle from AF2-based sampling and MD. Dashed lines indicate the extrema observed from 3DVA of the cryoEM dataset. Misfolded structures were omitted from analysis. **d)** Mb25 Nb-linker-HopQ angle from AF2-based sampling and MD. Dashed lines indicate the extrema observed from 3DVA of the cryoEM dataset. Misfolded structures were omitted from analysis. **e)** AF2-generated low quality structures from MSA subsampling at 16:32. **f)** SMO-PGS2 fusion MD simulations (left) and AF2 sampling (right).

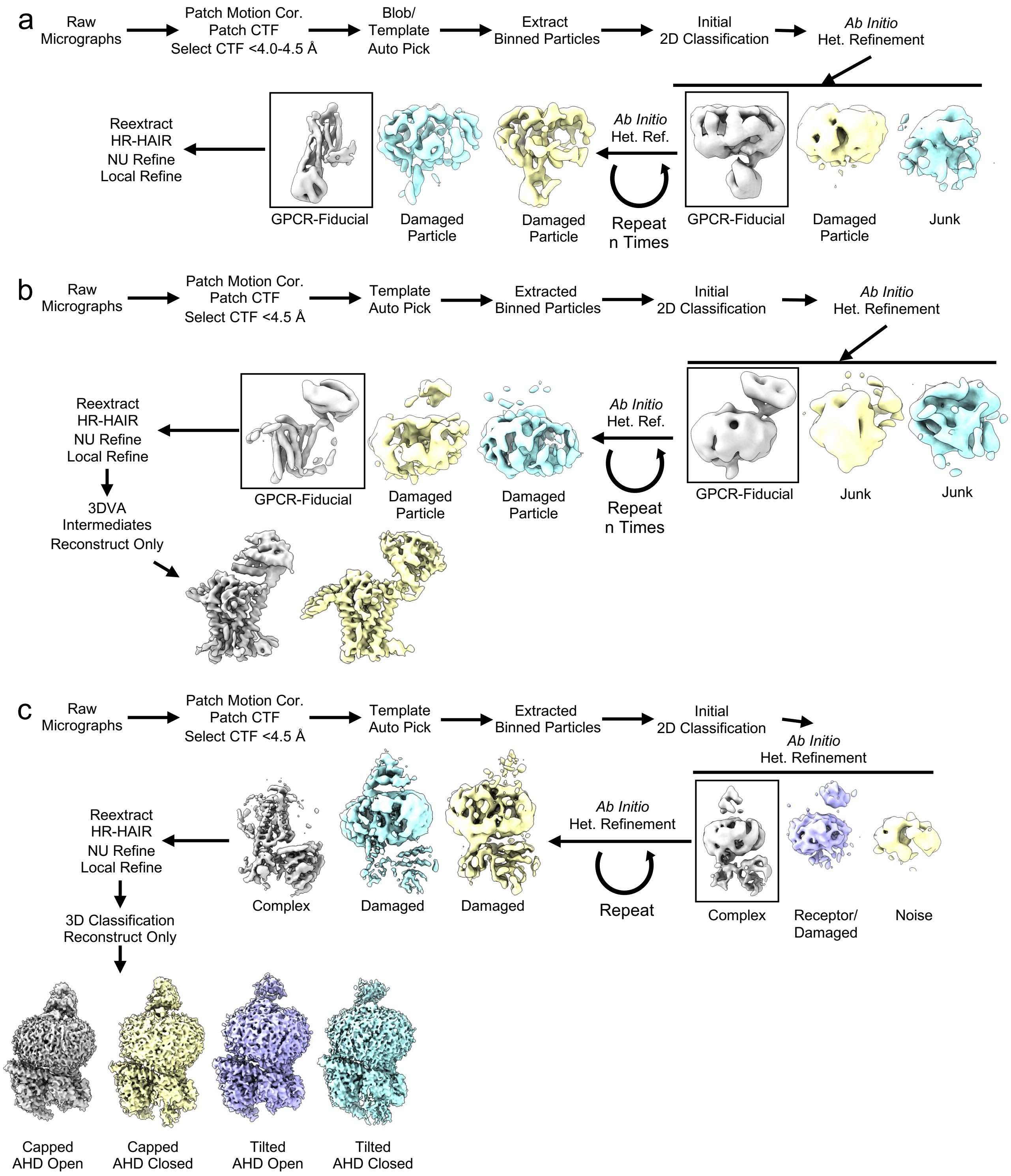


**Extended Data Figure 3: Cryo-EM data processing workflow a)** General processing workflow for the ICL fusion datasets **b)** General processing workflow for the BB3 fusion datasets **c)** Processing workflow for the BB3 fusion Gs complex dataset.

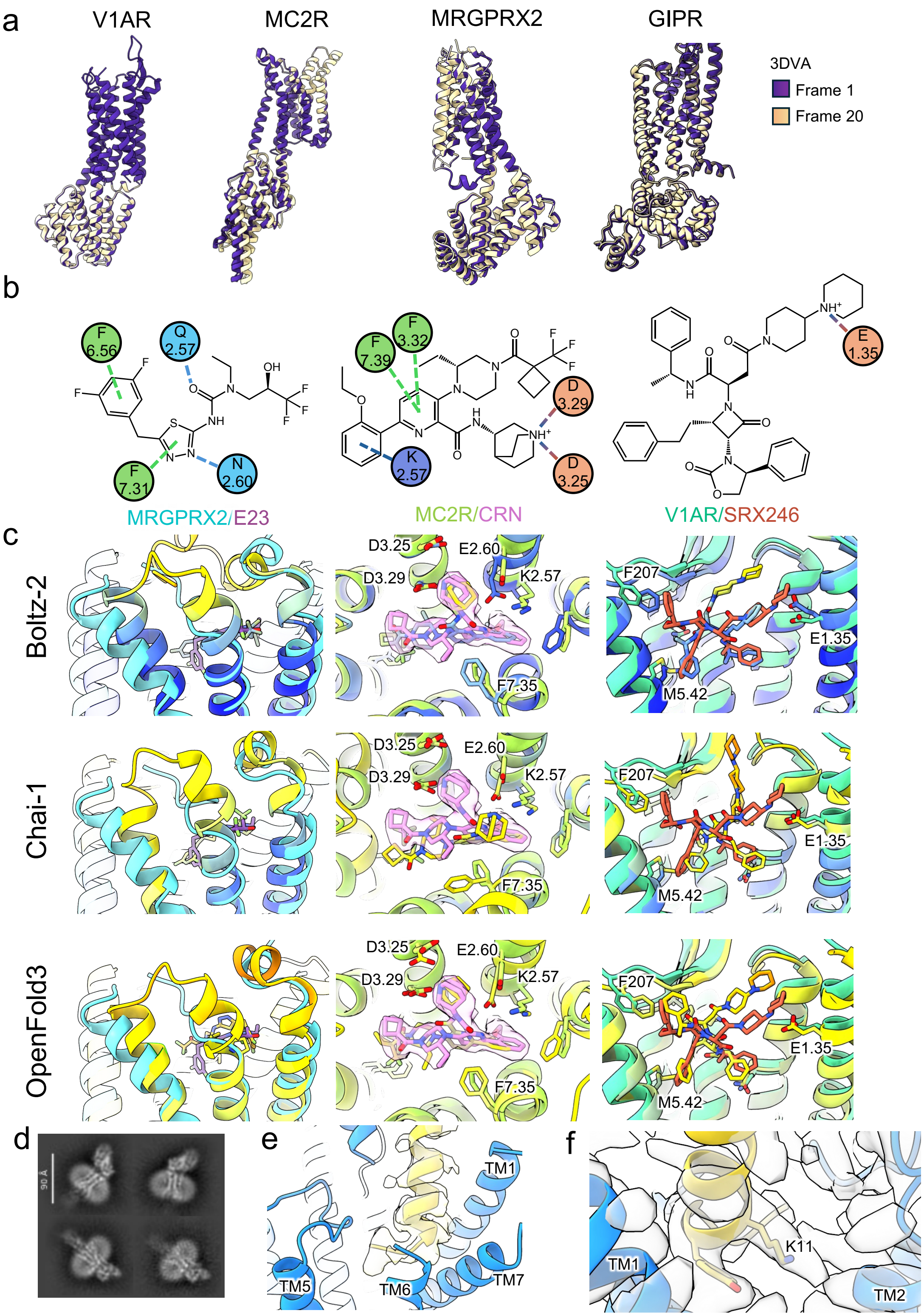


**Extended Data Figure 4: 3DVA, Co-folding, and Classification results for the ICL fusion structures determined in this work. a)** 3DVA first principal component for our four designs with our designs modeled into the first and last frame and aligned on the receptor transmembrane domain. **b)** Ligand interaction diagram for each ligand highlighting key interactions. **c)** Top predicted structure of each complex with Boltz-2, Chai-1, and OpenFold3 colored by pLDDT score and overlaid with the experimental structure. **d)** 2D Class averages of the GIPR consensus refinement. **e)** Map for GIP(5-31*) peptide (yellow) bound to GIPR (blue). **f)** Map showing the lipidated lysine of GIP(5-31*) (yellow) between TM1 and TM2 of GIPR (blue).

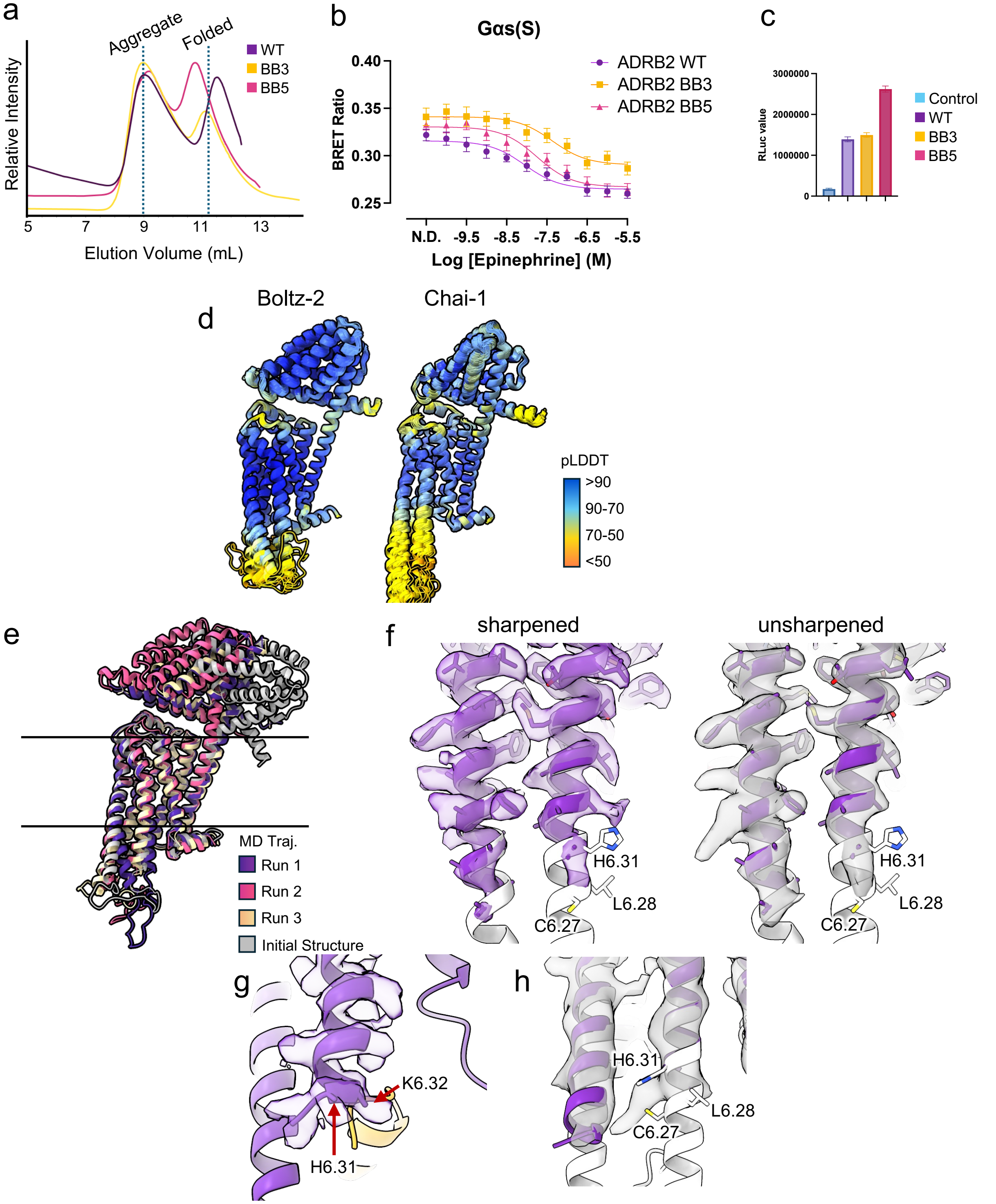


**Extended Data Figure 5: Biochemical, pharmacological, and cryoEM characterization of β2BB3 a)** Fluorescence size exclusion chromatography data for WT, BB3, and BB5 β2AR. **b)** TRUPATH BRET assays of WT, BB3, and BB5 β2AR signaling with epinephrine. N=3, n=4, data is mean +/- SEM. **c)** ELISA surface expression of WT, BB3, and BB5 β2AR. N=3, n=4, data is mean +/- SEM. **d)** Default OpenFold3, Boltz-2, and Chai-1 ensemble predictions of β2BB3 structures. **e)** MD simulation snapshot for β2BB3 starting from the tilted cryoEM structure (gray) in a lipid bilayer after 1 µs of simulation for three independent runs aligned on the receptor. **f)** Sharpened and unsharpened maps of the β2BB3-BI-167107 structure overlaid with the cryoEM structure as modeled (purple) and a hypothetical extended helical TM5 and TM6 (white) showing the location of labeling sites for smFRET/DEER. **g)** CryoEM map and model for β2BB3-BI-167107-Gs (purple, goldenrod) **h)** Overlay of the unsharpened map for β2BB3-propranolol with the modeled structure (purple) and and a hypothetical extended helical TM5 and TM6 (white) showing the location of labeling sites.

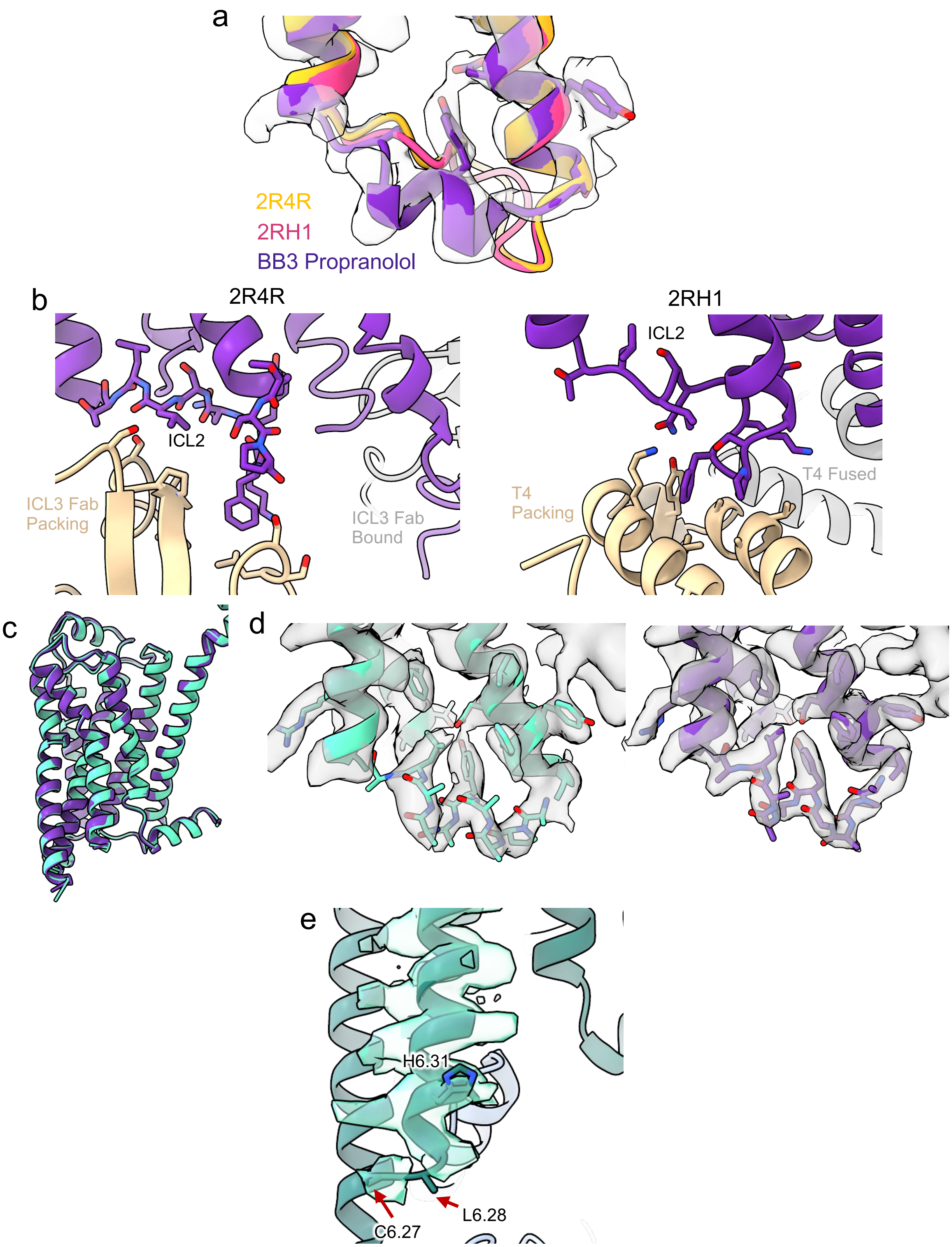


**Extended Data Figure 6: Comparison of key motifs for G-protein engagement in the β2BB3 structures a)** Overlay of ICL2 from the β2BB3-propranolol structure (purple) with the sharpened cryoEM map (gray) and crystal structures PDB:2R4R and 2RH1 of β2AR**. b)** Direct interactions of ICL2 from PDB:2R4R and 2RH1 with crystal packing neighbors. **c)** Overlay of the TM domain of cryoEM structures of β2BB3-BI-167107 (purple) and β2BB3-LM189 (teal). **d)** Sharpened cryoEM map (gray) and model for β2BB3-BI-167107 (purple) and β2BB3-LM189 (teal) highlighting the ICL2 region. **e)** CryoEM map and model for β2BB3-LM189-Gi (teal, slate blue PDB:9BUY).

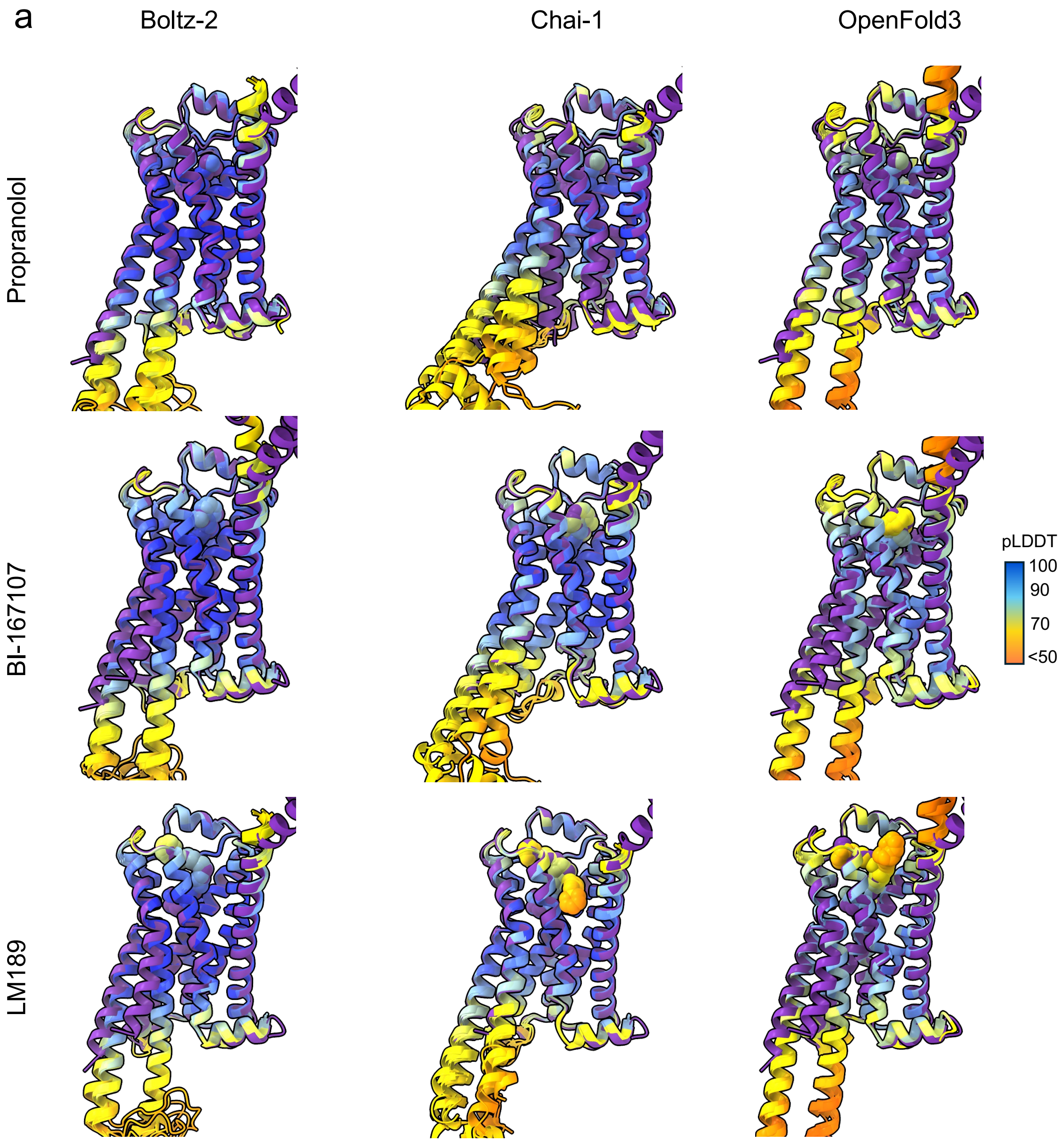


**Extended Data Figure 7: Comparison with Co-folding predictions for β2AR-ligand complexes a)** Overlay of cryoEM structures of β2BB3 bound to propranolol, BI-167107, and LM189 (purple) with all predictions from Boltz-2, Chai-1, and OpenFold3 colored by pLDDT score.
